# Language comprehension functionally modulates first-order relay thalamic nuclei

**DOI:** 10.1101/2025.10.30.685533

**Authors:** Liu Mengxing, Shiya Wang, Carmen Vidaurre, Sara Guediche, Garikoitz Lerma-Usabiaga, Pedro M. Paz-Alonso

## Abstract

Language, a uniquely human higher-order cognitive function, has traditionally been attributed to cortical mechanisms with limited attention given to subcortical contributions. Recent advances in non-invasive neuroimaging have revealed that thalamic activity can be modulated by attention and task demands. Moreover, lesion studies have hinted at the thalamus’s potential role in language processing. Nevertheless, the precise involvement of this structure in language remains unclear. Here, we ask whether language-related modulations can be detected as early as the sensory thalamic stage. Using functional MRI to image 40 human participants (both female and male) while processing linguistic and non-linguistic stimuli of three main language systems (reading, speech comprehension, and speech production), we demonstrate specific activation of first-order nuclei during targeted language system tasks: lateral geniculate (LGN) for reading, medial geniculate (MGN) for speech comprehension and ventrolateral (VLN) for speech production. Notably, we show linguistic versus non-linguistic stimuli exhibit functional modulations during comprehension tasks (reading and speech) of left MGN and, to a lesser extent, left LGN—in line with prior studies of lateralization of language processes. Multi-voxel classification analysis confirmed left-lateralized linguistic modulation in the MGN, but not in the LGN. Given the complexity of thalamic connectivity and its potential role in integrating sensory and cognitive processes, this work offers a comprehensive characterization of first-order thalamic engagement across reading, speech comprehension, and speech production within the same participants, advancing our understanding of thalamic involvement and thalamocortical interactions in language function.

## Introduction

Historically, neuroimaging research has unveiled the contributions of cortical regions to human language. The advent of sophisticated techniques such as functional magnetic resonance imaging (fMRI) has enabled researchers to explore the neural mechanisms underlying language functions in unprecedented spatial detail. This neuroimaging tool, combined with other techniques, have revealed that language processing engages a distributed network of cortical areas, each playing a unique role in aspects such as visual word form processing, phonological processing, syntactic analysis, and semantic comprehension (Binder & Desai, 2011; Friederici, 2011; Lau et al., 2008). While the role of the cerebral cortex in language is well-documented, the contributions of subcortical regions remain less explored, despite emerging evidence suggesting their importance (e.g., Abutalebi et al., 2013; Erb et al., 2013).

The classical cortico-centric perspective in cognitive neuroscience often neglects the intricate interplay between cortical and subcortical structures, which is essential for higher brain functions. Among subcortical structures, the thalamus has garnered attention for its role as a central hub (Shine et al., 2023), with reciprocal connections to virtually the entire cerebral cortex. Recent studies propose that the thalamus is integral to language processes, including speech production and comprehension (e.g., Barbas et al., 2013; Crosson, 2021; Fritsch et al., 2022). This shift in perspective underscores the need to reevaluate the subcortical contributions to language, particularly the thalamus, as a critical node in the language network. The thalamus receives orderly inputs from subcortical sensory, motor, and brainstem modulatory systems and it is massively interconnected with all areas of the ipsilateral neocortex in complex and highly specific patterns (Guillery & Sherman, 2002). Although first-order thalamic nuclei were historically characterized as straightforward relays transmitting sensorimotor information to cortex, it has long been recognized that even these nuclei are subject to top-down corticothalamic modulation rather than functioning as passive transponders (Mumford, 1991; Sherman & Guillery, 1996; Sillito et al., 1994). These nuclei, notably involved in receiving sensorimotor input from the periphery and transmitting it to specific cortical areas, have been foundational in shaping the classical understanding of thalamic function (Clascá, 2022), such as lateral geniculate nucleus (LGN) in visual system, medial geniculate nucleus (MGN) in auditory system and ventral lateral nucleus (VLN) in motor system.

More recently, however, research has revealed that many thalamic nuclei serve functions far beyond simple relay. These so-called higher-order thalamic nuclei do not merely transmit peripheral sensory signals but instead engage in complex cortico-thalamo-cortical interactions, facilitating information integration and coordination across distributed cortical networks (Sherman, 2007). For example, the mediodorsal (MD) thalamus plays a pivotal role in supporting prefrontal cortical functions, including cognitive flexibility, working memory, and decision-making (Mitchell, 2015). This emerging understanding reframes the thalamus as an active participant in shaping cortical processing rather than a passive intermediary, emphasizing its critical role in higher cognitive functions such as language, attention, and executive control. Notably, this non-relay function is not exclusive to higher-order thalamic nuclei; it extends to first-order nuclei like the LGN, MGN, and VLN. For example, the LGN can act as a node of cognitive control, with its responses modulated by attentional allocation (e.g., Ling et al., 2015; O’Connor et al., 2002; Schneider, 2011). The MGN likewise shows task-dependent modulation during speech recognition relative to non-speech control tasks (e.g., Mihai et al., 2021; von Kriegstein et al., 2008), demonstrating the broader functional versatility of the thalamus across both sensory and cognitive domains.

Reading, speech comprehension (SC) and speech production (SP) are some of the most relevant language systems used to convey most of the human communication that occurs daily. The sensorimotor features of these systems map on first-order relay thalamic nuclei: LGN, MGN, and VLN respectively. However, to date, neuroimaging research has not specifically examined the functional involvement and potential modulations of these thalamic nuclei in the same individual when they are engaging in language comprehension (i.e., reading and speech processing) and production (see Llano, 2013) for a review). Previous studies examining MGN involvement in SC revealed this nucleus exhibits heightened task-dependent activation during speech-recognition (e.g. syllable or speaker-identity judgments) tasks compared to control tasks (Diaz et al., 2012; Mihai et al., 2021; von Kriegstein et al., 2008). Note that these effects were obtained with active recognition paradigms (e.g., one-back syllable or speaker-identity judgements) rather than tasks requiring comprehension of lexical-semantic content. Despite these significant task-related findings concerning MGN involvement in different language tasks, we still do not know the extent to which first-order thalamic nuclei (related to different language systems) show functional modulations dependent upon whether the stimuli being processed are linguistic or non-linguistic.

Therefore, the objective of the present study is to investigate in the same individuals functional modulations of first-order thalamic nuclei (i.e., LGN, MGN, VLN). Specifically, we examine modulations associated with reading, SC, and SP depending on the linguistic nature of the stimuli. Essential to human communication, distinct language systems rely heavily on different types of sensorimotor information: reading relies on visual input, SC on auditory input, and SP on motor articulation. Here, we examine the engagement of first-order relay thalamic nuclei specifically in these three language systems: LGN in reading, MGN in SC, and VLN in SP. Our first hypothesis posits that first-order thalamic nuclei will exhibit modality-specific functional engagement in alignment with their relay function. Second, we hypothesize that their involvement will be modulated by the linguistic versus non-linguistic nature of the stimuli, and that this modulation will be left-lateralized. This prediction is strongest for the MGN, where task-dependent modulation during speech recognition tracks ability specifically in the left MGB (von Kriegstein et al., 2008; Mihai et al., 2019, 2021) and developmental dyslexia is associated with left MGB alterations (Diaz et al., 2012; Galaburda et al., 1994; Tschentscher et al., 2019). For the LGN, top-down modulation during visual processing is well established (Mumford, 1991; Sherman & Guillery, 1996; Sillito et al., 1994), whereas its left-lateralization is supported only by more indirect evidence from dyslexia (Giraldo-Chica et al., 2015; Müller-Axt et al., 2017, 2025; but see Díaz et al., 2018 for bilateral LGN modulation during visual speech); we therefore treat the latter as more exploratory. Third, we predict that functional coupling within each corresponding system (LGN–occipital cortex during reading, MGN–auditory cortex during SC, VLN–motor cortex during SP) will likewise be modulated by the linguistic nature of the stimuli. The present work builds in part upon the doctoral dissertation of Liu Mengxing (Mengxing, 2022).

## Methods

### Participants

The final study sample consisted of 40 right-handed native speakers of Basque (mean age 24.0 ± 4.6 years; 28 females). Data from additional five participants were excluded from further analyses due to excessive head motion during scanning. All participants had normal or corrected-to-normal vision, and none of them had a history of major medical, neurological, or psychiatric disorders. The study protocol was approved by the Ethics Committee of the Basque Center on Cognition, Brain and Language (BCBL). Prior to their inclusion in the study, all subjects provided informed written consent. Participants received monetary compensation for their participation.

### Design, materials, and experimental procedure

The functional MRI study employed a 3 x 2 within-subjects factorial block design with Modality (reading, SC, SP) and Stimuli (linguistic, non-linguistic) as factors. Linguistic tasks included single-word reading (visual), speech single-word comprehension (auditory), and single-word speech production (motor); while non-linguistic tasks involved viewing scrambled images (visual), listening to noise sequences (auditory), and producing unintelligible sounds (motor).

A total of 168 Basque noun words (e.g. *giltza*, *key* in English) were selected for the reading and SC modalities (average word frequency=13.9 [std=40.6], average length=6.4 letters [std=1.9 letters]). Half of the words were used in the reading modality in the form of visual images, while the remaining half were presented auditorily in the SC modality, with recordings made by a female Basque native speaker (no statistically significant differences between the two lists on word frequency and length, p=0.28 and p=0.87 respectively). For the non-linguistic reading stimuli, scrambled images were created by dividing the word images from the reading task into 10 × 10 pixel tiles and randomly shuffling them. For the non-linguistic SC stimuli, noise audios were generated with a custom MATLAB script that temporally randomized each spoken word by shuffling the samples of the audio waveform in a pseudo-random permutation (MATLAB randperm). This destroyed the phonological and semantic structure of the speech while preserving the duration, sampling rate, and total acoustic energy of the original stimulus. The stimulus-generation script is available in the study’s OSF repository (osf.io/kmhxz). These visual and auditory stimuli were counterbalanced across participants. For the linguistic SP stimuli, participants were asked to produce names of objects commonly found in their environment, such as “table” or “keyboard” corresponding to an office setting. For the non-linguistic motor stimuli, participants were instructed to produce unintelligible sounds, such as a palatal click, holding no semantic content. These unintelligible sounds were practiced with the participants prior to MRI scanning.

Participants completed seven functional runs, each comprising six 12-second activation blocks representing the experimental conditions: linguistic reading, linguistic SC, linguistic SP, non-linguistic reading, non-linguistic SC, and non-linguistic SP. Each activation block was separated by a 13.5 seconds rest interval to allow the BOLD signal to return to baseline. Data acquisition followed a sparse-sampling protocol (see *MRI data acquisition* section below), wherein each activation block contained 12 silence periods (1.2 seconds each) before each fMRI volume acquisition. The sparse-sampling protocol was designed to optimize the signal of interest for tasks involving auditory and motor modalities. To ensure comparability, it was applied to all three modalities.

During linguistic reading blocks, 12 single words were presented sequentially on the screen, and participants read them silently. In non-linguistic reading blocks, participants were instructed to fixate on 12 scrambled images based on selected words that appeared on the screen. Both blocks thus required the same directed fixation on centrally presented stimuli, minimizing systematic eye-movement differences between linguistic and non-linguistic blocks. For linguistic SC blocks, participants were prompted to close their eyes and listen to 12 spoken words. The non-linguistic SC blocks required participants to close their eyes and listen to 12 noise sequences. In linguistic SP blocks, participants produced one noun per silence period in the sparse-sampling sequence, with eyes closed. Similarly, in non-linguistic SP blocks, participants produced 12 unintelligible sounds, timed to each silence period. For all blocks where participants’ eyes were closed (i.e., SC and SP modalities across both linguistic and nonlinguistic stimuli), a bell sound was played at the end of each activation block, signaling participants to reopen their eyes.

### MRI data acquisition

Whole-brain MRI data acquisition was conducted on a 3-T Siemens PRISMA Fit whole-body MRI scanner (Siemens Medical Solutions) using a 64-channel whole-head coil. Functional images were acquired in a single gradient-echo echo-planar multiband pulse sequence with the following acquisition parameters: TE = 35 ms; MB acceleration factor = 5; 65 axial slices with a 2.4 mm^3^ voxel resolution; no inter-slice gap; flip angle = 56°; FoV = 210 mm; 1169 volumes in 7 runs. To optimize functional signals during tasks, we implemented a sparse-sampling paradigm (effective TR = 2.9 s, actual TR = 1.7 s), allowing a 1.2 s delay after each functional volume acquisition during which the scanner gradients were inactive, creating a silent MR environment. This silence period allowed for the presentation of acoustic or visual stimuli, or the performance of motor tasks without the acoustic scanner noise contamination (Hall et al., 1999). High-resolution structural images were collected for each participant using an MPRAGE T1-weighted sequence with parameters: TR = 2530 ms; TE = 2.36 ms; flip angle = 7°; FoV = 256 mm; voxel resolution = 1 mm^3^; 176 slices.

In total, 100 diffusion-weighted images were acquired using the anterior-to-posterior phase-encoding direction and 50 isotropically distributed diffusion-encoding gradient directions. The 100 diffusion-weighted images included 50 images with b-values of 1000 s/mm^2^ and 50 images with b-values of 2000 s/mm^2^. Twelve images with no diffusion-weighted values (b-values of 0 s/mm^2^) were obtained for motion correction and geometrical distortion correction, which comprised five images with the same phase-encoding direction as the DWI images and seven images with the reversed phase-encoding direction (posterior to anterior). Both DWIs and b0 images shared the following parameters: TR = 3600 ms, TE = 73 ms, FA = 78°, voxel size = 2 mm isotropic, 72 slices with no gap, and a multiband acceleration factor of 3.

### MRI data analysis

For structural image analysis, the T1w images were processed using RTP2 suit (Lerma-Usabiaga et al., 2023), which involves processing the subjects’ anatomical T1w image with recon-all from freesurfer (http://surfer.nmr.mgh.harvard.edu) and extract ROIs. First, Freesurfer was used to perform cortical/subcortical segmentation and parcellation. Next, the thalamic nuclei were obtained by running the thalamic segmentation module implemented in Freesurfer on a probabilistic atlas built based on histological and high-resolution *ex vivo* MRI data (Iglesias et al., 2018). For this study, we only considered left and right first-order relay nuclei as ROIs: the LGN, MGN and VLN. For functional and structural connectivity analyses, we also extracted the cortical regions V1/V2, A1 and M1. To parcellate the visual cortex we ran the Neuropythy (Benson & Winawer, 2018; https://github.com/noahbenson/neuropythy) tool on the Freesurfer results. A combination of the resulting V1 and V2 ROIs was used for our visual cortex ROI. A1 and M1 were converted from the human connectome project (HCP) atlas (Glasser et al., 2016). To convert them to individual subject space, we performed a non-linear registration of a 1 mm^3^ MNI template using Advanced Normalization Tools (ANTs, http://stnava.github.io/ANTs/).

For functional images preprocessing, we used SPM12 (Wellcome Center for Human Imaging, London) preprocessing routines and analysis methods. Images were corrected for differences in slice acquisition timing across every functional scan and then realigned for motion correction. Afterward, each subject’s functional volumes were smoothed using a 2 mm full-width half-maximum (FWHM) Gaussian kernel. Motion parameters were extracted from the realignment step to inform a volume repair procedure (ArtRepair; Stanford Psychiatric Neuroimaging Laboratory). This procedure identified unusable volumes based on scan-to-scan movement (>0.5 mm) and signal fluctuations in global intensity (>1.3%) and corrected for these volumes via interpolation from the nearest non-repaired scans. Five participants with more than 15% to-be-corrected outlier functional volumes were excluded. After volume repair, functional volumes were separately coregistered in two ways: 1) to MNI space to conduct whole-brain contrasts in normalized MNI space at the group level; 2) to high-resolution anatomical T1 images and resliced from the original 2.4 mm^3^ functional voxel dimensions to 1 mm^3^ voxels in anatomical T1 space for ROI analysis and functional connectivity. Finally, time series were temporally filtered to eliminate contamination from slow frequency drift (high-pass filter: 128 s).

**Statistical analyses** were performed on each subject data from both the MNI space and individual space using the general linear model (GLM). A series of impulses convolved with a canonical hemodynamic response function (HRF) were used to model the fMRI time series data. The six main experimental conditions in our design (i.e., 2 Task X 3 Modality) were modeled as epochs from the onset of the first trial to the last trial within the block. Given our sparse sampling design with an effective TR of 2.9s (TA = 1.7s, silent period = 1.2s), each block spanned 34.8s in real time (12 volumes × 2.9s). In our analysis, the epoch durations were specified as 20.4s, corresponding to the cumulative acquisition time (12 × 1.7s TA) within each block. These HRFs were used as covariates in the GLM. The motion parameters for translation (i.e., x, y, z) and rotation (i.e., yaw, pitch, roll) were used as covariates of no-interest in the GLM. SPM12 FAST was used for temporal autocorrelation modeling in this GLM due to its optimal performance removing residual autocorrelated noise in first-level analyses (Olszowy et al., 2019). The least-squares parameter estimates of the best-fitting canonical HRF’s height for each condition were used in pairwise contrasts. At the group level, whole-brain contrasts between conditions were computed in MNI space by performing paired t-tests on these images from different conditions, treating participants as a random effect. Three whole-brain contrasts were conducted: Reading - Others (SC + SP), SC - Others (Reading + SP) and SP - Others (Reading + SC). These three contrasts were selected to examine the modality-specific regions at the whole-brain level. Our standard statistical threshold for whole-brain maps was a p < 0.001 voxel-extent, Family Wise Error (FWE) corrected p < .05 at the cluster level.

**Individual ROI analysis** was performed on GLM results from the individual space data using the MARSBAR toolbox for its compatibility with SPM12. Given that this study focused on the involvement of first-order thalamic nuclei in language processing, the LGN, MGN and VLN were selected from both hemispheres as ROIs, defined in individual space using our thalamic segmentation implemented in FreeSurfer (Iglesias et al., 2018; https://freesurfer.net/fswiki/ThalamicNuclei). ROI activation and functional connectivity analyses were performed in each participant’s native anatomical space, using the native-space segmentations from the Iglesias et al.’s (2018) atlas, developed and validated in our group. We retained these rather than warping externally defined templates into native space, which would introduce registration errors that are particularly consequential for structures of this size. Relatedly, at our functional resolution (2.4 mm isotropic) the MGN ROI cannot be restricted to its first-order ventral subdivision, and contributions from higher-order subdivisions cannot be excluded; our findings therefore pertain to the MGN as a whole. Across participants, mean ROI volumes (± SD) were 276.0 ± 32.4 mm³ (left) and 263.0 ± 29.6 mm³ (right) for the LGN, 91.1 ± 11.9 and 100.1 ± 13.7 mm³ for the MGN, and 1457.7 ± 148.4 and 1454.5 ± 140.7 mm³ for the VLN. The LGN volumes are larger than those reported using dedicated high-resolution segmentation (≈121–132 mm³; Müller-Axt et al., 2017), while closer to the thalamic atlas proposed by Krauth et al. (2010) based on histological data (Left LGN: 232 and Right LGN: 224). This likely reflects differences in segmentation methods. The MGN volumes are comparable to prior estimates of whole-MGB volume (≈112–114 mm³; Mihai et al., 2019). After ROI definition, in line with our main experimental design, we extracted parameter estimates (i.e., scaled % signal change values; see (Mazaika, 2009) for each region and subject individually and used them as dependent variables in repeated-measures ANOVAs with subjects modeled as a within-subjects factor. Planned left-versus-right hemisphere comparisons were corrected for multiple comparisons using the Holm–Bonferroni procedure. One subject’s data was excluded due to technical failure of extracting ROI time series. To assess whether a dissociation of thalamic involvement exists between linguistic and non-linguistic tasks, we tested the left and right thalamic ROIs separately and compared their involvement between linguistic and non-linguistic tasks in their corresponding modality.

### Multi-voxel classification analysis

Prior to feature extraction and classification, all functional images were transformed to MNI space to be able to have the same number of voxels in each ROI and to perform the classification compiling trials of all subjects. Time series were extracted from preprocessed functional images described in the MRI data analysis section. The three first-order relay thalamic nuclei were selected, LGN, MGN and VLN -- left and right -- resulting in six ROIs. The number of voxels in each ROI was 29 and 28 in the left and right LGN respectively, 26 and 26 voxels in the left and right MGN, and 218 and 225 voxels in the left and the right VLN. Trends present in the time series were removed using a first-order polynomial regression in each run. Trials were obtained by averaging the time series starting 4s after the onset of each block and they were labeled using the corresponding stimulus. In total, we obtained 280 trials per class from all the subjects. Binary classification was performed between linguistic and non-linguistic tasks across reading, SC and SP modalities, using the associated thalamic ROIs (i.e. LGN, MGN, and VLN respectively). The separate classification using only the left or right thalamic ROIs was also performed to examine the linguistic-specific lateralization. A Linear Support Vector Machine (SVM) classifier with a soft margin and fixed random state (SVC function from Python scikit-learn with C=1.0, kernel=’linear’, random_state=42) was used due to i) its robustness to overfitting, ii) reduced sensitivity to outliers, and iii) strong performance with limited data. We employed eight-fold stratified cross-validation for train-test partitioning, with shuffling with a fixed random state applied at each iteration. Final results were reported using averaged Receiver Operating Characteristic (ROC) area under the curve (AUC) scores, in short AUC, evaluated on the test sets. Permutation tests with 10,000 iterations were then carried out to assess the significance of classification scores. Additionally, permutation tests for linguistic-specific lateralization were also performed. For that, in each iteration, we permuted the labels and performed the classification of the left and the right ROIs separately. Then, the AUC of the right side was subtracted from that of the left side. The difference was used as one sample in the reference distribution. The final permutation distribution consisted of 10,000 permuted samples, each representing the difference in classification performance between the left and right thalamic ROIs.

**Functional connectivity** was examined between each first-order thalamic nuclei of interest (LGN, MGN and VLN) and the site of cortical termination of the corresponding sensorimotor pathways (V1/V2, A1 and M1). The functional connectivity analyses were conducted using the beta-series correlation method (Rissman et al., 2004), implemented in SPM12 with custom Matlab scripts. We used beta-series correlation because our question concerned the magnitude of coupling between each nucleus and its corresponding primary cortex within a given condition, which this method estimates directly (Rissman et al., 2004). The canonical HRF in SPM was fit to each trial within each experimental condition and the resulting parameter estimates (i.e., beta values) were sorted according to the study conditions to produce a condition-specific beta series for each voxel. Pairwise functional connectivity analysis between thalamic nuclei and cortical regions was conducted at the individual-subject level for linguistic and non-linguistic stimuli in the corresponding sensorimotor modality (LGN-V1V2 in Reading, MGN-A1 in SC and VLN-M1 in SP). The beta-series correlation values (r values) were transformed to Fisher’s z values by applying an arc hyperbolic tangent transform (Fisher, 1915) at the subject level for each pair of ROIs and each experimental condition. Since the correlation coefficient is inherently restricted to the range from −1 to +1, this transformation ensured the null hypothesis sampling distribution approached that of the normal distribution. To assess the significance of the correlation findings at the group level, the z-transformed correlation of the individual subjects was compared against zero at the group level. We also tested linguistic versus non-linguistic differences using simple-effects comparisons on the subject-level beta-series z-transformed values.

**Structural connectivity** analysis was conducted by using the tractography protocol proposed by Liu et al (2022) to reconstruct the first-order relay thalamic tracts on the DWI data collected in this study. Three pairs of first-order thalamic tracts were reconstructed: bilateral optic radiations (OR), acoustic radiations (AR), and motor radiations (MR). The FA profile of each tract from each subject was obtained using the RTP2- pipeline (Lerma-Usabiaga et al., 2023). The mean of the FA profile was calculated as the index by averaging the 100 FA values in the FA profile for each tract and each subject. To examine the associations between structural connectivity and regional activation, correlation analyses were conducted between the FA values from the structural connectivity and the percent signal change from the thalamic nuclei for each first-order thalamocortical pathway of interest (corresponding to each modality). This same analysis was also conducted between the FA values from structural connectivity and the left *versus* right difference in percent signal change for each thalamic nuclei corresponding to each modality task.

## Results

### Whole-brain contrasts

To identify brain regions associated with processing specific modalities across all participants and linguistic *versus* non-linguistic stimuli, we computed three specific whole-brain contrasts: i) Reading > Others (SC + SP); SC > Others (Reading + SP) and SP > Others (Reading + SC). These contrasts revealed the involvement of the first-order thalamic relay nuclei and the primary corresponding sensorimotor cortical regions in modality-specific tasks. The Reading > Others contrast showed increased activation in bilateral LGN and primary and secondary visual cortex (V1/V2) during reading compared to when participants process and produce speech (Figure 1*A*). Similarly, the SC > Others contrast revealed engagement of bilateral MGN and primary auditory cortex (A1; Figure 1*B*). Finally, the SP > Others contrast showed higher involvement of the motor thalamic nuclei (bilateral VLN) and primary motor cortex (M1). Vermis III in the cerebellum also showed specific involvement in the motor contrast (Figure 1*C*). Using an inclusive mask of the thalamus with these contrasts revealed the specificity of each of them showing functional activation of the expected thalamic nuclei: LGN for Reading > Others contrast; MGN for SC > Others contrast; and, VLN extended to mediodorsal nucleus for the SP > Others contrast (Figure 1*D*).

**Figure 1.**
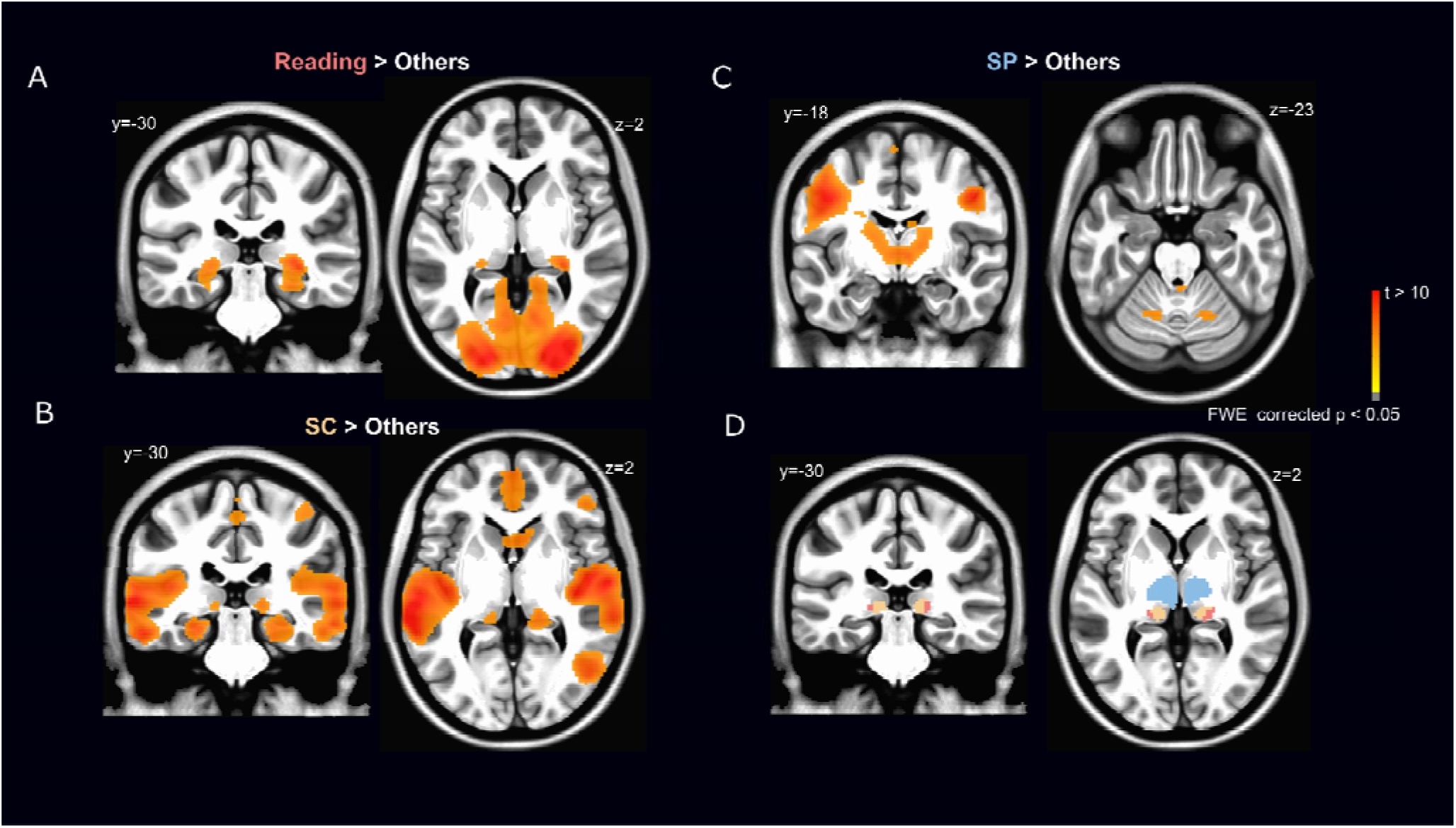
Brain sections showing whole-brain activations across all subjects for the A) Reading > Others (SC +SP) contrast, B) SC > Others (Reading + SP) contrast, C) SP > Others (Reading + SC) contrast. D) Brain sections showing the same three contrasts using an inclusive whole-thalamus mask (coral = Reading > Others, gold = SC > Others, blue = SP > Others). All brain sections presented here are in MNI space. The statistical threshold was set to p < 0.001 voxel-extent, FWE-corrected < .05 at the cluster level.

### ROI results

ROI analyses were conducted to characterize the activation profile of the three thalamic ROIs (LGN, MGN and VLN) bilaterally for the main experimental stimuli in the three sensorimotor modalities. We extracted fMRI parameter estimates from these six ROIs and conducted hypothesis-driven analyses based on planned comparisons between conditions. In line with our hypotheses, first we examined the specificity of the engagement of each sensorimotor thalamic nuclei per each of the three language-related tasks, collapsing across Hemisphere and Stimuli factors. For that, we conducted three separated one-way ANOVAs per each thalamic nuclei (i.e., LGN, MGN, VLN) examining the factor Modality (i.e., Reading, SC, SP).

The one-way ANOVA for the LGN revealed a statistically significant main effect of Modality (*F*(2,76) = 30.0; *p* < 0.001, = 0.44). Post-hoc Holm-corrected paired comparisons indicated this was due to a significantly higher percentage signal change (PSC) for Reading compared to both SC (mean difference = 0.57, 95% CI [0.24, 0.90], p < 0.001, d_z_=0.65) and SP (mean difference = 1.39, 95% CI [1.06, 1.72], p < 0.001, d_z_=1.07) (see Figure 2). Additionally, SC showed significantly higher PSC than SP (mean difference= 0.81, 95% CI [0.49, 1.15], *p* < 0.001, d_z_=0.70). Similarly, the one-way ANOVA for the MGN also revealed a significant main effect of Modality (*F*(2,76) = 4.6; *p* = 0.014, = 0.11). Post-hoc Holm-corrected paired comparisons showed that SC had significantly higher PSC compared to Reading (mean difference = 0.85, 95% CI [0.19, 1.52], *p* < 0.01, d_z_=0.73) and SP (mean difference = 1.16, 95% CI [0.50, 1.83], *p* < 0.05, d_z_=0.42). Finally, for the VLN, the one-way ANOVA also revealed a significant main effect of Modality (*F*(2,76) = 42.3; *p* < 0.001, = 0.53). Post-hoc Holm-corrected paired comparisons indicated that SP elicited significantly higher PSC than both Reading (mean difference = 1.54, 95% CI [1.27, 1.81], *p* < 0.001, d_z_=1.17) and SC (mean difference = 1.21, 95% CI [0.94, 1.49], *p* < 0.001, d_z_=0.93). A significant difference was also found between Reading and SC modalities (mean difference = 0.33, 95% CI [0.05, 0.60], *p* < 0.05, d_z_=0.72).

**Figure 2.**
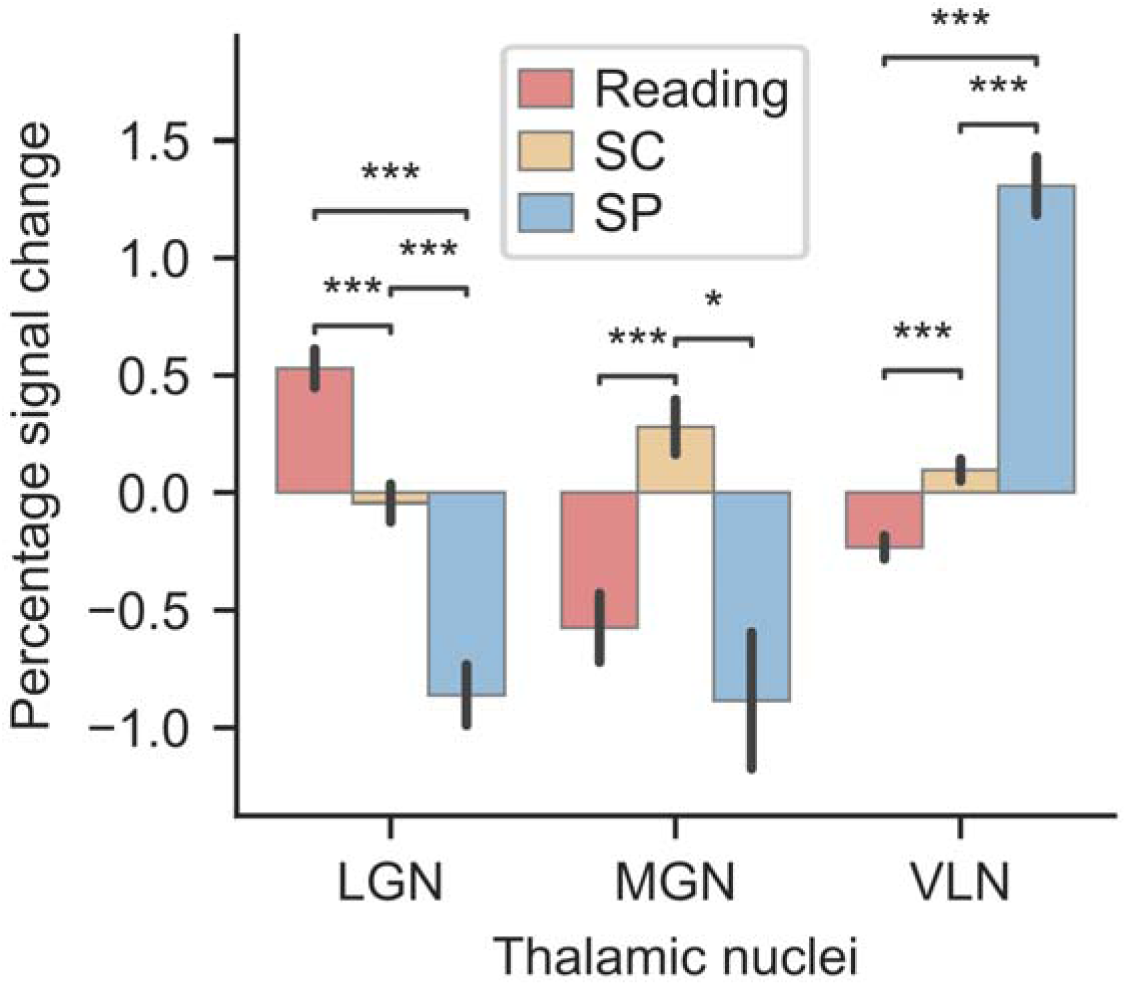
ROI analysis. Scaled percentage signal change for bilateral LGN, MGN and VLN first-order relay thalamic nuclei as a function of sensorimotor Modality: Reading, SC and SP. Error bars represent the standard error mean. *p < 0.05, **p < 0.01, ***p < 0.001.

Second, in line with our predictions, we examined whether the linguistic nature of the stimuli modulated the recruitment of first-order thalamic nuclei as a function of lateralization. To this end, we conducted three separate Stimuli × Hemisphere repeated-measures ANOVAs for each thalamic nucleus (LGN, MGN, VLN), with Stimuli (linguistic, non-linguistic) and Hemisphere (left, right) as factors. The Stimuli main effect was significant in the LGN (F(1,38) = 4.44, p = 0.042, ^2^ = 0.105), reflecting an overall linguistic advantage (linguistic > non-linguistic across both hemispheres), but not in the MGN (F(1,38) = 1.92, p = 0.174, ^2^ = 0.048) or VLN (F(1,38) = 2.33, p = 0.136, ^2^ = 0.058). The Stimuli × Hemisphere ANOVAs revealed a significant interaction in the MGN (F(1,38)=8.63, p=0.006, ^2^ =0.19), but not in the LGN (F(1,38) = 1.02, *p* = 0.318, η²p = 0.026) or VLN (F(1,38) = 0.52, p = 0.474, η²p = 0.014). Given the significant MGN interaction, we decomposed it into simple effects of Stimuli within each hemisphere; for the LGN, we examined the left-hemisphere comparison as a directional, theoretically motivated test (see Introduction). First, we examined the effect of stimuli (reading words vs. scrambled images) on the LGN using paired t-tests. The results revealed a significant difference between linguistic and non-linguistic conditions in the left LGN (t = 2.43, p = 0.02, d_z_=0.39), but not in the right LGN (t = 1.41, p = 0.18, d_z_=0.23). Next, we conducted the same t-tests for the MGN, comparing responses to listening to speech sounds vs. scrambled sounds. Again, a significant difference was found in the left MGN (t = 3.00, p < 0.01, d_z_=0.48), while no significant effect was observed in the right MGN (t = ™0.27, p = 0.78, d_z_=-0.04). Finally, we examined the effect of stimuli (speech production vs. unintelligible speech production) on the VLN. No significant differences were found for either left VLN (t = −1.59, p = 0.12, d_z_ = −0.26) or right VLN (t = −1.37, p = 0.18, d_z_ = −0.22).

### Multi-voxel classification results

On top of ROI analysis based on activation amplitude, we further tested whether the activation pattern of the ROIs differ between linguistic and nonlinguistic tasks using multi-voxel classification analysis. First, we conducted separate classification on LGN, MGN and VLN, to classify linguistic *versus* nonlinguistic tasks within their corresponding modalities (Figure 3B). The results showed that neither left and right LGN activation classify significantly linguistic *versus* nonlinguistic reading tasks (*AUC* = 0.54, *p* = 0.12, and *AUC* = 0.52 *p* = 0.23, respectively); left MGN (*AUC* = 0.59, *p* = 0.001), but not right MGN (*AUC* = 0.50, *p* = 0.47), classify significantly linguistic *versus* nonlinguistic speech comprehension tasks; and, both left and right VLN significantly classify linguistic *versus* nonlinguistic in SP tasks (left VLN *AUC* = 0.73, *p* < 0.001, and right VLN *AUC* = 0.73, *p* < 0.001). To examine lateralization in terms of multi-pattern classification, we tested whether the classification score was significantly higher in left *versus* right MGN and VLN, since these were the only nuclei showing linguistic versus non-linguistic classification (Figure 3C). The results revealed that the left MGN has a statistically significant higher classification score than the right MGN (*AUC* difference = 0.08, significant at the 97.5th percentile), while this lateralization effect was not observed in the VLN (*AUC* difference = 0, *p* > 0.05).

**Figure 3.**
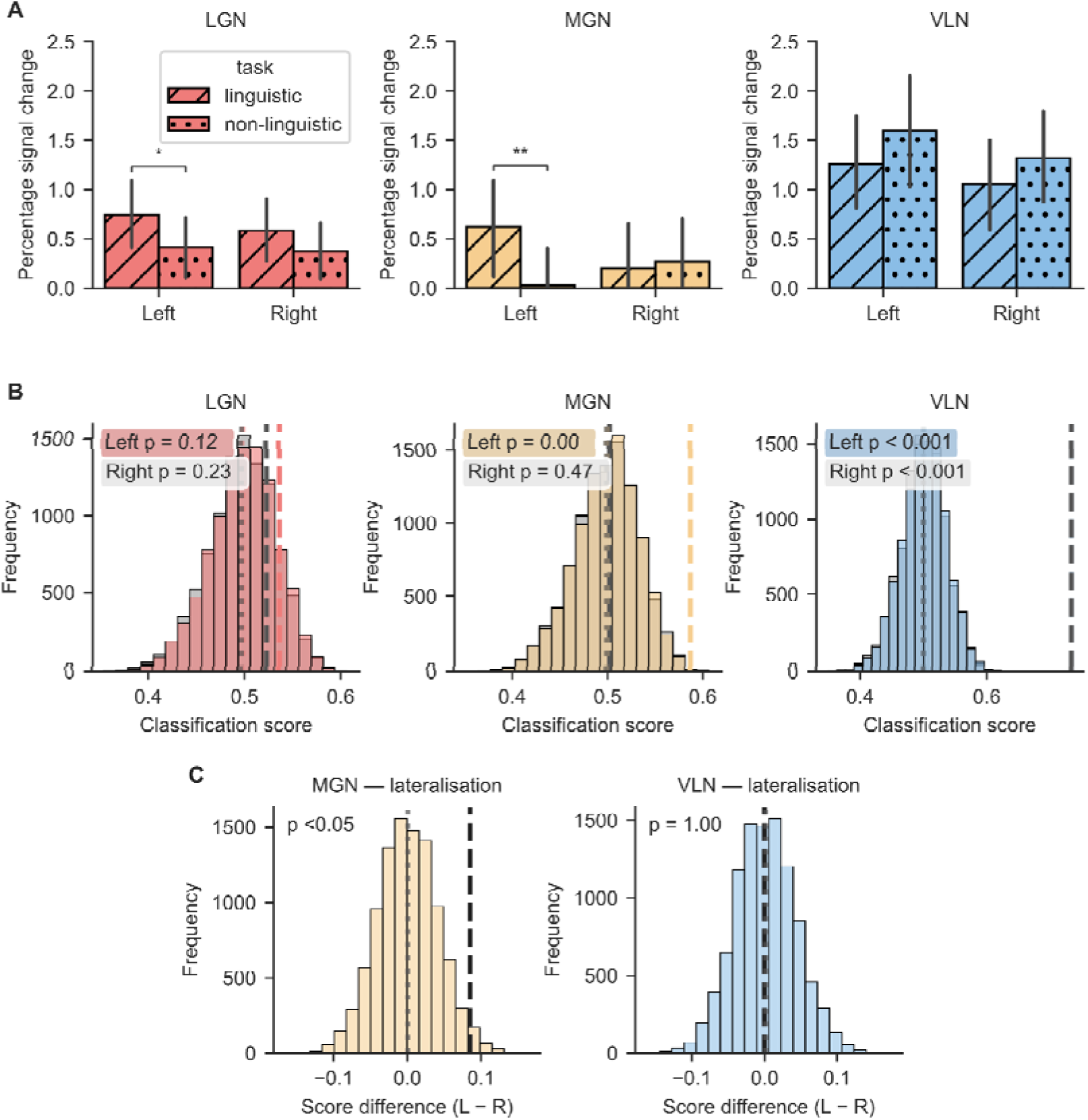
**A.** ROI analysis. Scaled percentage signal change for the LGN, MGN, and VLN first-order relay thalamic nuclei as a function of Hemisphere and Stimuli. Error bars represent the standard error mean. *p < 0.05. **B.** Classification analysis. Histograms showing distributions of 10,000 permutation scores for the left and right hemisphere. Dashed vertical lines indicate real classification scores. Dotted vertical lines show empirical chance levels derived from permutation means. P-values indicate the proportion of permutations that exceeded the real classification performance. **C.** Lateralization of classification. Histograms showing distributions of classification score difference between left and right (Left - Right scores) from 10,000 permutations. Red dashed lines indicate real lateralization differences. Blue dotted lines show empirical chance levels (mean of permutation differences). P-values are two-tailed tests indicating whether observed lateralization differs significantly from chance-level asymmetry.

### Functional and structural connectivity results

Pairwise functional connectivity analyses of three first-order relay thalamic pathways for the linguistic and non-linguistic tasks of the corresponding modalities were examined: LGN-V1/V2 visual pathway for linguistic and non-linguistic tasks in the visual modality; MGN-A1 auditory pathway for linguistic and non-linguistic tasks in the auditory modality; and, VLN-M1 motor pathway for linguistic and non-linguistic tasks in the motor modality. To examine the significance of the correlation findings at the group level, Fisher’s z-transformed correlation values of individual subjects were compared against zero. The results revealed that predicted functional connections between regions based on known neuroanatomy were statistically significant for both linguistic (Figure 4A) and non-linguistic tasks (Figure 4B). In the visual pathway, LGN-V1/V2 functional coupling for linguistic reading (z = 0.56, *p* < 0.001 for left and z = 0.58, *p* < 0.001 for right) and non-linguistic reading tasks (z = 0.52, *p* < 0.001 for left; z = 0.51, *p* < 0.001 for right) was statistically significant. Similarly significant functional coactivation was observed between MGN and A1 for linguistic and non-linguistic speech processing tasks, as well as between the VLN and M1 for motor linguistic and non-linguistic speech production tasks. Simple-effect analyses were conducted to examine if there were statistically significant differences between the functional connectivity in linguistic versus non-linguistic tasks for each of these first-order thalamic pathways. None of the comparisons revealed statistically significant differences (*p*s ≥ 0.54).

**Figure 4.**
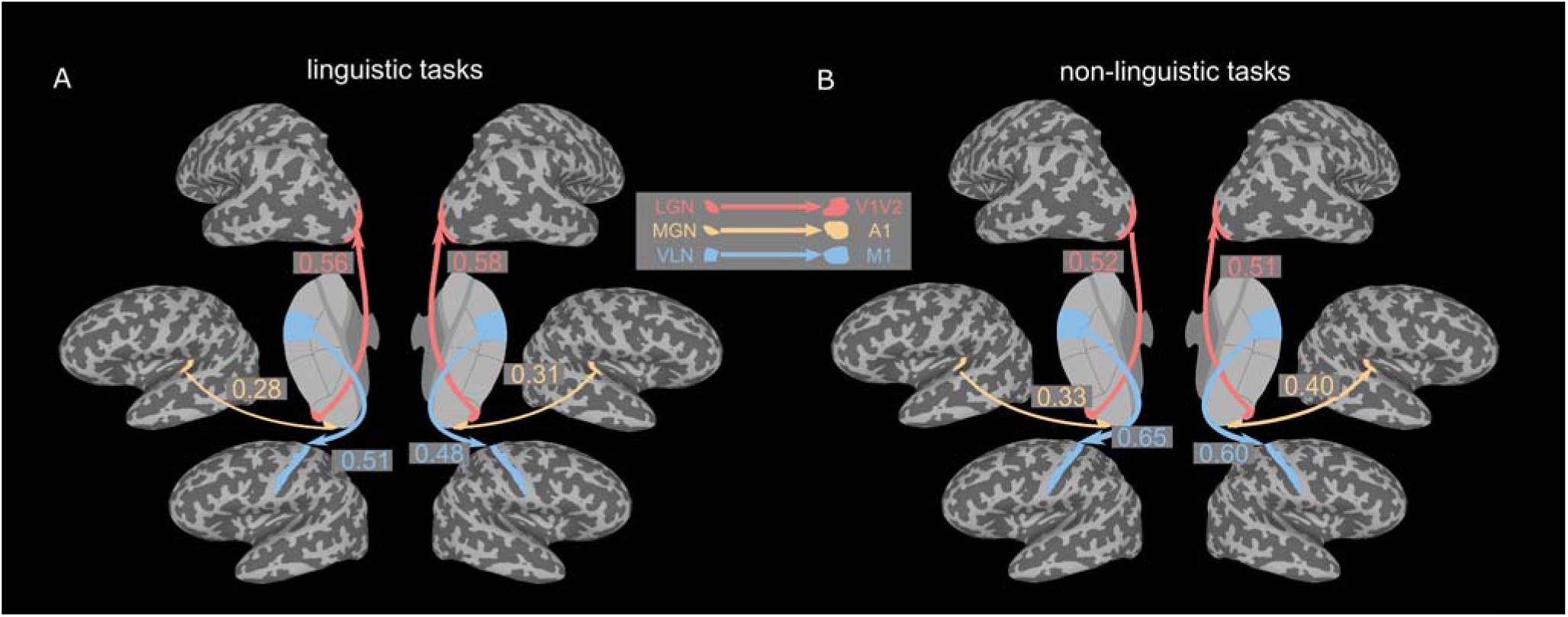
Functional connectivity analysis of three first-order relay thalamic pathways in A) linguistic and B) non- linguistic tasks for each of the corresponding modalities. Lines with an arrow represent the connection of the first-order thalamic relay nuclei with the cortical primary sensorimotor regions. Thalamic nuclei, cortical regions and the lines are colored as a function of sensorimotor modality: coral indicates visual modality, gold indicates auditory modality and blue indicates motor modality. The values along the arrow lines are the Fisher’s-transformed z values. All the reported z values are significant against zero with p < 0.001 after Bonferroni FWE correction for multiple comparisons.

To examine the association between structural connectivity and regional activation, we reconstructed three pairs of tracts bilaterally: the OR, the AR and the MR. FA values were extracted as an index of white matter integrity for each tract. Correlation analyses were conducted between the FA values of each of the left and right white-matter tracts corresponding to each modality task (i.e., OR for reading, AR for speech comprehension, and MR for speech production) and the respective percent signal change of left and right LGN, MGN and VLN linguistic and non-linguistic tasks in each modality. Also, differences in percent signal change in left versus right for the LGN and MGN were correlated to left and right OR and AR as well as for the differences in FA between left and right OR and AR. None of the regression analyses were statistically significant (*p*s _≥_ 0.08).

## Discussion

The present study was aimed at investigating functional modulations of first-order thalamic nuclei (i.e., LGN, MGN, VLN) during reading, speech comprehension, and language production depending on the linguistic nature of the stimuli. Our findings revealed specific engagement of the LGN, MGN and VLN for linguistic and non-linguistic tasks in their corresponding modality (i.e., LGN for reading, MGN for speech comprehension, VLN for language production). Importantly, linguistic *versus* non-linguistic tasks showed left-lateralized modulations of the LGN for reading and the MGN for speech comprehension tasks. Functional coupling within each thalamocortical network (e.g., LGN-V1) was statistically significant and also specific to the corresponding language modality (e.g., reading). These results are discussed next.

### Language systems as first-order thalamic nuclei localizers

In the present study, we aimed to identify how the reliance of the main language systems on visual aspects (as in reading), auditory aspects (as in speech comprehension), and motor aspects (as in speech production) can be used to functionally localize three main first-order relay nuclei individually: LGN, MGN, and VLN. That was the case in the group analyses shown in Figure 1, which map the main thalamocortical networks involved in each of the three language systems, and was also evident at the single-subject level. Aside from a few exceptions (e.g., Crosson, 2021; Mihai et al., 2019, 2021), most of the neuroimaging research in the neurobiology of language has focused on examinations of the cortical network involved in language-related processes. Indeed, many highly relevant models of the neurobiology of language have been put forward. Nevertheless, even though the thalamocortical network identified here for reading, speech comprehension, and speech production are a common signature and are present in neuroimaging studies examining these language systems, these thalamocortical circuitries have not been mapped in detail, nor have they been incorporated into mainstream neurobiology of language models (Friederici, 2011; Hickok & Poeppel, 2007).

While practically any modality-specific task can localize first-order thalamic nuclei (e.g., O’Connor et al., 2002), our language-related tasks offer the advantage of simultaneously localizing these nuclei and characterizing their engagement during language processing. By leveraging the inherent sensory demands of reading, SC, and SP, we successfully mapped the functional engagement of the LGN, MGN, and VLN, respectively. The strong modality-specific activation observed in these nuclei (and their corresponding cortical counterparts) underscores their role as first-order relay stations that selectively process language-related sensorimotor information. This approach is particularly valuable for studies investigating thalamocortical language networks, as it ensures that anatomical boundaries are defined under the same cognitive conditions as the primary language contrasts of interest.

We note that, beyond these first-order nuclei, our results also revealed mediodorsal (MD) thalamus activation during speech production (Figure 1). This finding suggests the MD may contribute to higher-order cognitive aspects of the motor planning or memory retrieval involved in language production (Crosson, 1985; Mitchell, 2015; Pergola et al., 2018; Stockert et al., 2023). There are two possible explanations for this observation: (1) unlike the online processing demands of reading and speech comprehension, speech production in our experiment required participants to remember and produce words and sounds that were practiced outside the scanner. Given that the MD thalamus is a critical hub for memory function, particularly in retrieval and executive control of learned sequences, its engagement in speech production may reflect these functions (Mitchell & Chakraborty, 2013; Pergola et al., 2018); (2) Given that speech production is a complex motor behavior requiring coordination between phonological, articulatory, and motor planning processes, MD activation may reflect its role in modulating prefrontal-motor interactions necessary for fluent verbal output. This notion aligns with previous research demonstrating that lesions in the MD nucleus can result in disfluencies and impaired verbal initiation, further emphasizing its importance in language-related motor execution (Crosson, 1985; Stockert et al., 2023).

### Left-lateralized linguistic modulation of LGN and MGN during language comprehension

A key finding of our study is that linguistic processing induces a left-lateralized modulation of LGN and MGN activity. Specifically, we observed that compared to non-linguistic stimuli, processing linguistic stimuli in the visual and auditory modalities selectively enhanced activation in LGN and MGN, respectively. Previous research has demonstrated that LGN (O’Connor et al., 2002; Schneider, 2011) and MGN (Diaz et al., 2012; Mihai et al., 2021) activation can be modulated by task demands and attention. Crucially, we found that this linguistic modulation is left-lateralized, reinforcing the well-established left-hemisphere dominance for language (Binder et al., 1995; Frost et al., 1999). For example, Binder et al. (1995) reported that while processing non-speech stimuli for pitch and simple sequence information activated bilateral auditory cortices, semantic processing tasks elicited left-lateralized activity across distributed language-related cortical regions. Our findings extend this cortical asymmetry to the thalamus, demonstrating that language-related lateralization is not confined to cortical areas but is evident at early sensory processing stages within the thalamus itself. The fact that this modulation is left-lateralized also argues against a purely attentional account, since a generic difference in engagement between conditions would not predict a specifically lateralized response; attentional allocation is, in any case, in part intrinsic to linguistic processing rather than a separable confound.

While both the LGN and MGN exhibited left-lateralized activation for linguistic tasks in amplitude-based ROI analyses, the left MGN additionally showed significant multi-voxel discrimination, but not the right MGN, suggesting more fine-grained linguistic encoding in the left auditory thalamus. This result is consistent with the MGN role in spectrotemporal speech analysis (see von Kriegstein et al., 2008), where left-lateralized phonological processing may drive both amplitude and spatial pattern differences. On the one hand, in line with previous findings (e.g., O’Connor et al., 2002), the observed LGN amplitude modulation suggests top-down linguistic influences on visual processing. On the other hand, the lack of significant discrimination between linguistic and non-linguistic tasks in the LGN may reflect either methodological limitations in detecting fine-grained activity patterns in this region or that these top-down effects manifest through global signal enhancement rather than precise visual word form encoding within the LGN itself.

Unlike the LGN and MGN, the VLN did not show a modulatory engagement for the production of speech relative to the production of unintelligible sounds. We speculate that this is due to the greater specificity of the motor circuits compared to auditory and visual circuits. One of the first structures that has been related to vocalization was the face area of the motor cortex, which is located in the lateral part of the precentral cortex in primates. The main input structure of the motor cortex is the VLN, and one of the main input structures of the VLN is the cerebellum. Non-human primate studies have shown that a large number of cells in the VLN activate before vocalization onset (Farley, 1997). Thus, the initial phase of vocal preparation may be happening in the motor circuits before reaching the VLN, so this motor nuclei is simply relaying cerebellar information to the motor cortex. This may explain why we did not observe differences between speech production and producing unintelligible sounds in VLN in the present study.

### Functional and structural connectivity did not reveal linguistic modulations

Our results showed significant functional coupling between first-order relay thalamic nuclei and their corresponding cortical counterparts within both left and right hemispheres. This finding reveals that the expected functional coactivation was present across our experimental conditions. In relation to structural connectivity, no associations were observed between functional indexes, including regional activation of left and right thalamic nuclei, and the difference in activation among them. Previous studies showing first-order thalamic nuclei attentional or language-related modulations typically found these effects at the regional level and, as far as we know, no previous MRI studies have specifically examined whether or not these modulations can be also observed in functional coupling and in association with structural connectivity. To our knowledge, few studies have examined modulatory effects on the functional and structural connectivity levels, specifically using a widely utilized procedure to test functional connectivity in task-related fMRI and state-of-the-art tractography analyses. However, the absence of observed linguistic modulations in terms of functional coactivation does not necessarily imply that these modulations cannot be found using other functional and/or structural connectivity approaches, such as effective connectivity (e.g., Dynamic Causal Modelling, Structural Equation Modelling) or other diffusion MRI approaches. Future studies should explore other connectivity approaches to confirm or expand upon the results reported here.

The present study is not without limitations. Firstly, due to scanner noise, the auditory modality is typically more susceptible to scanner noise contamination compared to the other modalities. Although we optimized our experimental procedures and employed a sparse-sampling approach to minimize the influence of scanner noise across all language modalities, the inherent challenge of dissociating speech-related neural activity from scanner noise in auditory tasks remained a limitation. For instance, future mitigation of this issue can be achieved by incorporating advanced real-time acoustic denoising techniques. Another limitation of this study concerns the stimuli selection used to assess linguistic modulations. Linguistic effects were identified by comparing real word processing with nonsensical stimuli in both visual and auditory modalities. However, word processing exists on a continuum, and intermediate stimulus types, such as pseudowords and consonant strings, may engage thalamic nuclei differently. Linguistic and non-linguistic stimuli also differ in low-level sensory properties (e.g., spatial frequency in the visual modality, spectrotemporal statistics in the auditory modality) to which these nuclei are sensitive, so a low-level contribution to the present effects cannot be fully excluded. That said, a purely low-level account would not readily predict the left-lateralized pattern we observed, which is more consistent with language-related lateralized processing. Nevertheless, further examination is needed to determine whether or not the effects from these intermediate linguistic stimuli are similar or vary on a continuum, and can provide a deeper understanding of the mechanisms underlying the thalamic linguistic modulation observed in this study.

## Conclusion

Using an optimized experimental design, including sparse-sampling to minimize effects of scanner noise, this study successfully established a novel functional localization framework for sensorimotor thalamic nuclei in linguistic tasks. Notably, this study characterized the language-specific involvement of first-order thalamic nuclei in language processing, with left-lateralized linguistic modulation observed in LGN and MGN, reinforcing the well-documented left-hemisphere dominance for language and extending it to the early thalamic processing of visual and auditory linguistic input. Future research should further investigate the precise mechanism of language-specific modulation in sensory thalamus.

## Acknowledgement and Funding

The authors thank Jake Vinnacombe for proofreading and helpful comments on the manuscript. L.M. was supported by grants from the European Union’s Horizon 2020 research and innovation programme under the Marie Sklodowska-Curie (grant agreement No. 713673) and from “la Caixa” Foundation (grant no. 11660016 and grant no. 100010434 under the agreement no. HR18-00178- DYSTHAL). G. L.-U. was supported by the Spanish Ministry of Science and Innovation (PID2021-123577NA-I00) and Basque Government (PIBA_2022_1_0014). P.M.P-A. was supported by grants from the Spanish Ministry of Science and Innovation (PID2021-123574NB-I00), from the Basque Government (PIBA-2021-1-0003), and from “la Caixa” Foundation (ID 100010434) under the agreement HR18-00178-DYSTHAL. BCBL acknowledges funding from the Basque Government through the BERC 2022–2025 program and from the Spanish State Research Agency through the BCBL Severo Ochoa excellence accreditation CEX2020-001010/AEI/https://doi.org/10.13039/501100011033.

## Notes

Conflict of interest: The authors declare no competing financial interests.

### Competing Interest Statement

The authors have declared no competing interest.

### Summary of Updates

Figures revised; ROI analysis statistics revised, effect size provided. Language revised.

## References

Abutalebi, J., Rosa, P. A. D., Castro Gonzaga, A. K., Keim, R., Costa, A., & Perani, D. (2013). The role of the left putamen in multilingual language production. Brain and Language, 125(3), 307–315. 10.1016/j.bandl.2012.03.009

Barbas, H., García-Cabezas, M. Á., & Zikopoulos, B. (2013). Frontal-thalamic circuits associated with language. Brain and Language, 126(1), 49–61. 10.1016/j.bandl.2012.10.001

Benson, N. C., & Winawer, J. (2018). Bayesian analysis of retinotopic maps. eLife, 7, 1–29. 10.7554/eLife.40224

Binder, J. R., & Desai, R. H. (2011). The neurobiology of semantic memory. Trends Cogn Sci, 15(11), 527–536. 10.1016/j.tics.2011.10.001

Binder, J. R., Rao, S. M., Hammeke, T. A., Frost, J. A., Bandettini, P. A., Jesmanowicz, A., & Hyde, J. S. (1995). Lateralized Human Brain Language Systems Demonstrated by Task Subtraction Functional Magnetic Resonance Imaging. Archives of Neurology, 52(6), 593–601. 10.1001/archneur.1995.00540300067015

Clascá, F. (2022). A Brief History of Thalamus Research. In The Thalamus (pp. 1–26). Oxford University Press.

Crosson, B. (1985). Subcortical functions in language: A working model. Brain and Language, 25(2), 257–292. 10.1016/0093-934X(85)90085-9

Crosson, B. (2021). The Role of Cortico-Thalamo-Cortical Circuits in Language: Recurrent Circuits Revisited. Neuropsychology Review, 31(3), 516–533. 10.1007/s11065-019-09421-8

Díaz, B., Blank, H., & von Kriegstein, K. (2018). Task-dependent modulation of the visual sensory thalamus assists visual-speech recognition. Neuroimage, 178, 721–734. 10.1016/j.neuroimage.2018.05.032

Diaz, B., Hintz, F., Kiebel, S. J., & von Kriegstein, K. (2012). Dysfunction of the auditory thalamus in developmental dyslexia. Proceedings of the National Academy of Sciences, 109(34), 13841–13846. 10.1073/pnas.1119828109

Erb, J., Henry, M. J., Eisner, F., & Obleser, J. (2013). The brain dynamics of rapid perceptual adaptation to adverse listening conditions. Journal of Neuroscience, 33(26), 10688–10697.

Farley, G. R. (1997). Neural firing in ventrolateral thalamic nucleus during conditioned vocal behavior in cats. Experimental Brain Research, 115(3), 493–506. 10.1007/PL00005719

Fisher, R. A. (1915). Frequency Distribution of the Values of the Correlation Coefficient in Samples from an Indefinitely Large Population. Biometrika, 10(4), 507–521. 10.2307/2331838

Friederici, A. D. (2011). The brain basis of language processing: From structure to function. Physiological Reviews, 91(4), 1357–1392. 10.1152/physrev.00006.2011

Fritsch, M., Rangus, I., & Nolte, C. H. (2022). Thalamic Aphasia: A Review. Current Neurology and Neuroscience Reports, 22(12), 855–865. 10.1007/s11910-022-01242-2

Frost, J. A., Binder, J. R., Springer, J. A., Hammeke, T. A., Bellgowan, P. S. F., Rao, S. M., & Cox, R. W. (1999). Language processing is strongly left lateralized in both sexes. Evidence from functional MRI. Brain, 122(2), 199–208. 10.1093/brain/122.2.199

Galaburda, A. M., Menard, M. T., & Rosen, G. D. (1994). Evidence for aberrant auditory anatomy in developmental dyslexia. Proceedings of the National Academy of Sciences, 91(17), 8010–8013. 10.1073/pnas.91.17.8010

Giraldo-Chica, M., Hegarty, J. P., & Schneider, K. A. (2015). Morphological differences in the lateral geniculate nucleus associated with dyslexia. NeuroImage. Clinical, 7, 830–836. 10.1016/j.nicl.2015.03.011

Glasser, M. F., Coalson, T. S., Robinson, E. C., Hacker, C. D., Harwell, J., Yacoub, E., Ugurbil, K., Andersson, J., Beckmann, C. F., Jenkinson, M., Smith, S. M., & Van Essen, D. C. (2016). A multi-modal parcellation of human cerebral cortex. Nature, 536(7615), 171–178. 10.1038/nature18933

Guillery, R. W., & Sherman, S. M. (2002). Thalamic relay functions and their role in corticocortical communication: Generalizations from the visual system. Neuron, 33(2), 163–175. 10.1016/S0896-6273(01)00582-7

Hall, D. A., Haggard, M. P., Akeroyd, M. A., Palmer, A. R., Summerfield, A. Q., Elliott, M. R., Gurney, E. M., & Bowtell, R. W. (1999). “Sparse” temporal sampling in auditory fMRI. Human Brain Mapping, 7(3), 213–223. 10.1002/(sici)1097-0193(1999)7:3%3C213::aid-hbm5%3E3.0.co;2-n

Hickok, G., & Poeppel, D. (2007). The cortical organization of speech processing. Nat Rev Neurosci, 8(5), 393–402. 10.1038/nrn2113

Iglesias, J. E., Insausti, R., Lerma-Usabiaga, G., Bocchetta, M., Van Leemput, K., Greve, D. N., van der Kouwe, A., Fischl, B., Caballero-Gaudes, C., & Paz-Alonso, P. M. (2018). A probabilistic atlas of the human thalamic nuclei combining ex vivo MRI and histology. NeuroImage, 183(July), 314–326. 10.1016/j.neuroimage.2018.08.012

Krauth, A., Blanc, R., Poveda, A., Jeanmonod, D., Morel, A., & Székely, G. (2010). A mean three-dimensional atlas of the human thalamus: Generation from multiple histological data. NeuroImage, 49(3), 2053–2062. 10.1016/j.neuroimage.2009.10.042

Lau, E. F., Phillips, C., & Poeppel, D. (2008). A cortical network for semantics: (De)constructing the N400. Nature Reviews Neuroscience, 9(12), 920–933. Doi 10.1038/Nrn2532

Lerma-Usabiaga, G., Liu, M., Paz-Alonso, P. M., & Wandell, B. A. (2023). Reproducible Tract Profiles 2 (RTP2) suite, from diffusion MRI acquisition to clinical practice and research. Scientific Reports, 13(1), Article 1. 10.1038/s41598-023-32924-7

Ling, S., Pratte, M. S., & Tong, F. (2015). Attention alters orientation processing in the human lateral geniculate nucleus. Nature Neuroscience, 18(4), 496–498. 10.1038/nn.3967

Llano, D. A. (2013). Functional imaging of the thalamus in language. Brain and Language, 126(1), 62–72. 10.1016/j.bandl.2012.06.004

Mazaika, P. (2009). Percent Signal Change for fMRI calculations. Manual, (1), 1–8.

Mihai, P. G., Moerel, M., de Martino, F., Trampel, R., Kiebel, S., & von Kriegstein, K. (2019). Modulation of tonotopic ventral medial geniculate body is behaviorally relevant for speech recognition. eLife, 8, e44837. 10.7554/eLife.44837

Mihai, P. G., Tschentscher, N., & von Kriegstein, K. (2021). Modulation of the Primary Auditory Thalamus When Recognizing Speech with Background Noise. The Journal of Neuroscience, 41(33), 7136–7147. 10.1523/jneurosci.2902-20.2021

Mitchell, A. S. (2015). The mediodorsal thalamus as a higher order thalamic relay nucleus important for learning and decision-making. Neuroscience and Biobehavioral Reviews, 54, 76–88. 10.1016/j.neubiorev.2015.03.001

Mitchell, A. S., & Chakraborty, S. (2013). What does the mediodorsal thalamus do? Frontiers in Systems Neuroscience, 7. 10.3389/fnsys.2013.00037

Müller-Axt, C., Anwander, A., & von Kriegstein, K. (2017). Altered Structural Connectivity of the Left Visual Thalamus in Developmental Dyslexia. Current Biology, 27(23), 3692–3698.e4. 10.1016/j.cub.2017.10.034

Müller-Axt, C., Kauffmann, L., Eichner, C., & von Kriegstein, K. (2025). Dysfunction of the magnocellular subdivision of the visual thalamus in developmental dyslexia. Brain, 148(1), 252–261. 10.1093/brain/awae235

Mumford, D. (1991). On the computational architecture of the neocortex. I. The role of the thalamo-cortical loop. Biological Cybernetics, 65(2), 135–145. 10.1007/BF00202389

O’Connor, D. H., Fukui, M. M., Pinsk, M. A., & Kastner, S. (2002). Attention modulates responses in the human lateral geniculate nucleus. Nature Neuroscience, 5(11), 1203–1209. 10.1038/nn957

Olszowy, W., Aston, J., Rua, C., & Williams, G. B. (2019). Accurate autocorrelation modeling substantially improves fMRI reliability. Nature Communications, 10(1), Article 1. 10.1038/s41467-019-09230-w

Pergola, G., Danet, L., Pitel, A. L., Carlesimo, G. A., Segobin, S., Pariente, J., Suchan, B., Mitchell, A. S., & Barbeau, E. J. (2018). The Regulatory Role of the Human Mediodorsal Thalamus. Trends in Cognitive Sciences, 22(11), 1011–1025. 10.1016/j.tics.2018.08.006

Rissman, J., Gazzaley, A., & D’Esposito, M. (2004). Measuring functional connectivity during distinct stages of a cognitive task. NeuroImage, 23(2), 752–763. 10.1016/j.neuroimage.2004.06.035

Schneider, K. A. (2011). Subcortical Mechanisms of Feature-Based Attention. Journal of Neuroscience, 31(23), 8643–8653. 10.1523/JNEUROSCI.6274-10.2011

Sherman, S. M. (2007). The thalamus is more than just a relay. Current Opinion in Neurobiology, 17(4), 417–422. 10.1016/j.conb.2007.07.003

Sherman, S. M., & Guillery, R. W. (1996). Functional organization of thalamocortical relays. Journal of Neurophysiology, 76(3), 1367–1395. 10.1152/jn.1996.76.3.1367

Shine, J. M., Lewis, L. D., Garrett, D. D., & Hwang, K. (2023). The impact of the human thalamus on brain-wide information processing. Nature Reviews Neuroscience, 1–15. 10.1038/s41583-023-00701-0

Sillito, A. M., Jones, H. E., Gerstein, G. L., & West, D. C. (1994). Feature-linked synchronization of thalamic relay cell firing induced by feedback from the visual cortex. Nature, 369(6480), 479–482. 10.1038/369479a0

Stockert, A., Hormig-Rauber, S., Wawrzyniak, M., Klingbeil, J., Schneider, H. R., Pirlich, M., Schob, S., Hoffmann, K.-T., & Saur, D. (2023). Involvement of Thalamocortical Networks in Patients With Poststroke Thalamic Aphasia. Neurology, 100(5), e485–e496. 10.1212/WNL.0000000000201488

Tschentscher, N., Ruisinger, A., Blank, H., Díaz, B., & von Kriegstein, K. (2019). Reduced structural connectivity between left auditory thalamus and the motion-sensitive planum temporale in developmental dyslexia. Journal of Neuroscience, 39(9), 1720–1732. 10.1523/JNEUROSCI.1435-18.2018

von Kriegstein, K., Patterson, R. D., & Griffiths, T. D. (2008). Task-Dependent Modulation of Medial Geniculate Body Is Behaviorally Relevant for Speech Recognition. Current Biology, 18(23), 1855–1859. 10.1016/j.cub.2008.10.052

